# Paleometabolomics reveals impacts of abiotic factors on rodent midden metabolism over the last 50,000 years

**DOI:** 10.64898/2026.02.26.708167

**Authors:** Thomas Dussarrat, Francisca P. Díaz, Victoria Gödde, Marcus Persicke, Cédric Cassan, Claudia Rouveyrol, Karsten Niehaus, Pierre Pétriacq, Caroline Müller, Claudio Latorre, Rodrigo A. Gutiérrez

## Abstract

Metabolomics and paleoecology combined can reveal how past ecosystems worked, helping us predict future changes more accurately. Pioneering studies are needed to shed light on the potential of the so-called paleometabolomics and to standardise its application. Here, we deployed an untargeted metabolomic workflow on a timeline (200 to 49,600 cal yr BP) of rodent middens that efficiently and reproducibly captured rodent midden metabolic diversity, recovering 79% of the richness detected in 15 contemporary plant species. We found that midden chemical diversity and metabolites were influenced by the midden composition, age, and environment. Variation at the metabolite level in middens could fit age, past temperature and precipitation levels with an *R²* > 88% and their plant composition. Compounds and families responding to climate variation included lipids (*e.g.* glycerophospholipids) and other metabolites linked to redox status such as phenolics (*e.g.* flavonoids, lignans). The responses of significant midden chemical indices and compounds to abiotic pressures were supported by their response in plants collected near the midden sites to temperature and soil water content across an elevation gradient. Overall, our results not only showcase paleometabolomics as a powerful tool to reconstruct past ecosystem dynamics and metabolic evolutionary trajectories, but also to uncover relevant chemical families that could serve as trackers of past -and potentially future- climate fluctuations.

## 1. Introduction

Paleoecology seeks to explore and understand the past ecological dynamics of the biosphere, providing long-term context for present-day ecosystems and their future trajectories (Jackson & Williams 2004). Paleoecological and historical records can be used to explore past community structures and their responses to major climate events, and reveal, for instance, lost biotic connections to support restoration programs (Bush *et al*. 2022; Wood *et al*. 2013). Paleogenomics, the analysis of ancient DNA, where “ancient” refers to samples over 100 years old (Brunson & Reich 2019), has revolutionised this research field (Díaz *et al*. 2019; Napier *et al*. 2020). Paleometabolomics, the comprehensive analysis of all metabolites in ancient samples, may have considerable potential and provide complementary information beyond that provided by genomics (Bromage *et al*. 2026).

Metabolomics is currently advancing rapidly and has earned considerable interest in diverse research fields such as ecology (Walker *et al*. 2022) and biogeosciences (Abs *et al*. 2025). The integration of metabolomics and paleoecology only depends on the emergence of a methodology developed by a few pioneering studies (Badillo-Sanchez *et al*. 2023; Bromage *et al*. 2026). The use of paleometabolomics is supported by the properties of metabolism and chemicals. First, although variable, certain chemicals such as lipids, pigments or terpenes can persist for millions of years (Glass *et al*. 2012; Marynowski *et al*. 2007; Summons *et al*. 2022). In contrast, DNA recovery can be extended to the early Pleistocene (hundreds of thousands of years) (Dalén *et al*. 2023). Organic material and archaeological artefacts can preserve metabolic information for thousands of years (Badillo-Sanchez *et al*. 2023; Crown *et al*. 2015; Velsko *et al*. 2017). Second, metabolism has proven to be extremely effective at capturing environmental variations (Defossez *et al*. 2021; Dussarrat *et al*. 2022). Metabolomics has also been useful in unravelling chemical aspects of ecological processes (Díaz *et al*. 2024; Müller & Junker 2022). Hence, paleometabolomics opens remarkable possibilities, from reconstructing past human behaviour (Brockbals *et al*. 2018; Cole *et al*. 2025; Zimmermann *et al*. 2021) to refining ecological models and providing guidance for crop breeding through records of chemical adaptations and ecosystem responses to major climate events (Meiri & Bar-Oz 2024; Wood *et al*. 2013).

Paleometabolomics may be subjected to similar limitations as paleogenomics. For instance, the susceptibility of metabolites to degradation or contamination from exogenous compounds, which result from the interaction with the environment, could challenge the interpretation of the findings. In fact, the turnover varies considerably between metabolites, where some (e.g. fructose-bisphosphate) are highly reactive while others (e.g. lipids, terpenes, phenolics) are stable over ages. However, despite postdepositional alteration, ancient samples retained a chemical profile comparable to fresh tissues (Brownstein *et al*. 2020; Velsko *et al*. 2017). Besides, the analysis of the variation in richness and diversity of chemical families could complement studies at the metabolite level, where residual enzyme activities may affect concentrations (Defossez *et al*. 2021; Dussarrat *et al*. 2025; Müller & Junker 2022). Moreover, the question of whether metabolic regulations within ancient samples compare with those in recent samples remains unclear (Jackson & Williams 2004). Thus, establishing baselines for the models using recent tissues and supporting findings with observations on recent material is required to ensure that insights from the past relate to opportunities tomorrow (Bush & McInerney 2013; De Porras *et al*. 2015; Jackson & Williams 2004).

The use of omics techniques on ancient plant samples is scarce (Brunson & Reich 2019), especially to study the response of plant communities to climatic events. Herbaria have offered important opportunities for paleogenomics, yet their use has been limited due to restricted resource availability and a lack of detailed geographical and historical context (Burbano & Gutaker 2023; Eckert *et al*. 2025). In contrast, rodent middens consist of plant and animal remains preserved within a matrix of crystallised rodent urine and are typically found in protected rock crevices and shelters (Díaz *et al*. 2019; Latorre *et al*. 2003). Rodent middens were screened to provide a reliable spatiotemporal record of plant and animal communities (Lesser & Jackson 2011; Nowak *et al*. 2000). For instance, middens were used to reconstruct plant communities using plant macrofossils, pollen microfossils or ancient DNA (aDNA) (De Porras *et al*. 2015; Díaz *et al*. 2019). Besides, fecal pellet size accurately predicted late Quaternary precipitation dynamics (González-Pinilla *et al*. 2021) and *n*-alkane composition of middens was linked to past climate change in the Atacama Desert (Frugone-Álvarez *et al*. 2023). Middens were found across diverse environments spanning elevational, latitudinal, and climatic gradients, offering the opportunity to apply consistent methodologies across sites. Importantly, these records also include samples of different ages, providing a multi-temporal archive that potentially captures 50,000 years of ecological change (Becklin *et al*. 2024; Latorre *et al*. 2003). Overall, rodent middens emerged as an ideal tool for reconstructing past metabolic strategies adopted by plants or other organisms to face climatic threats (Becklin *et al*. 2024; Díaz *et al*. 2019).

Here, we used a temporally discontinuous timeline of rodent middens up to 49,600 years from the Atacama Desert to investigate (i) whether untargeted metabolomics can be used to reproducibly capture ancient metabolism and (ii) how the influence of age and climate variation on rodent midden chemistry is characterised. We hypothesised that major chemical classes such as phenolics and fatty acids could be used to track past climate change (Dussarrat *et al*. 2022). More specifically, we used technical replicates of middens to assess the reproducibility of our method to characterise primary and specialised metabolites of rodent middens at both metabolite and chemical family levels. We then linked variation in metabolic intensities or richness and diversity of major chemical families to midden composition (*i.e.* number of species and phylogeny), age, temperature and precipitation anomalies as well as CO_2_ levels. To support the responses of midden chemicals to abiotic pressures, we assessed the response of these chemicals to temperature and precipitation levels in fifteen recent plant species collected across an elevation gradient near the midden sites.

## 2. Materials and Methods

### 2.1. Plant and midden samples

Fossil rodent middens were previously collected from the Antofagasta Region of northern Chile. These paleoenvironmental archives include records from the Cerros de Aiquina (CDA) site, which spans an elevational range between 3,100 and 3,380 m.a.s.l., and a single record from the Vegas de Tilocalar (VDT), located at 2,414 m.a.s.l. (Díaz *et al*. 2019; Wood *et al*. 2019). At each collection site, modern local vegetation within an approximate foraging radius of 50 m was characterised and documented to provide an ecological baseline (Díaz et al. 2012; Becklin et al. 2024). Indurated midden samples were extracted from the host rock using flooring chisels and hammers (A Holmgren *et al*. 2023). To prevent contamination by modern biological material, the outer weathered surface layer was carefully removed during collection. The remaining material was transported to the laboratory and processed using a multi-proxy approach (A Holmgren *et al*. 2023), including subsamples for radiocarbon dating, plant macrofossil identification, pollen analysis, and aDNA metabarcoding. A dedicated subsample was additionally allocated for paleometabolomics analyses, which constitute the focus of the results presented in this study.

Plant samples were previously collected in analogue locations as the midden samples (Talabre-Leja transect, 22-24°S), along an elevation gradient (Dussarrat *et al*. 2022). The aerial parts of the 15 species used in this study were harvested on April 6 and 7, 2019, at one to six elevation levels with at least three biological replicates per elevation, as described previously (Dussarrat *et al*. 2022). Samples were snap frozen in liquid nitrogen in the Atacama Desert and transported in dry ice to the laboratory, where they were stored until freeze-drying for 48 h. From the freeze-dried powders, an equal amount (10 mg) from the same quantity of replicates for each altitude was pooled in a tube for each plant species (*e.g.* for *Atriplex imbricata*, a mix of three replicates collected at 3,570, 3,370, 3,270 and 2,870 m.a.s.l).

### 2.2. Metadata collection

Middens were radiocarbon dated using accelerator mass spectrometry (Latorre *et al*. 2002, 2003). Radiocarbon dates were calibrated using the Southern Hemisphere ShCal13 calibration curve as previously described (Díaz *et al*. 2019). For each midden, subsamples were extracted (*i.e.* collecting the midden core) under sterile conditions to extract aDNA, which was sequenced using Illumina’s next-generation pair-end sequencing technology (MiSeq, Reagent Kit v3) to identify the relative abundance and identity of plant species contained in the sample (Díaz *et al*. 2019). Species composition was not available for four middens: CDA_595B, CDA_571, CDA_563B, and CDA_593A. For these samples, species composition was approximated based on the plant species currently present at the collection site (Table S1). This strategy was supported by the fact that rodents are dietary generalists and that midden composition reflects the plant diversity of their environment (Becklin *et al*. 2024). Phylogenetic diversity (PD) was calculated using Faith’s index (Faith 1992) based on a presence–absence matrix and a pruned phylogenetic tree obtained from the Open Tree of Life via the *rotl* package in R (v.4.5.1) (Michonneau *et al*. 2016). The *pd()* function from the *picante* package (Kembel *et al*. 2010) was used, assigning equal branch lengths when branch-length information was not provided for the retrieved phylogeny. To place midden-derived biodiversity patterns within a broader climatic context, we compiled paleoclimate metadata describing temperature anomalies and carbon dioxide (CO_2_) levels in Antarctica over the last 800,000 years (Jouzel *et al*. 2007; Lüthi *et al*. 2008). A high correlation between temperature anomalies and CO_2_ levels was observed (Lüthi *et al*. 2008). Precipitation anomalies represent the difference between the estimated mean annual rainfall (MAR), which considers the elevation and location of the midden, and the modern MAR (González-Pinilla *et al*. 2021). However, precipitation anomalies were not available for four of the middens (Table S1). Metadata for plant samples, which include elevation, temperature and soil water content, were obtained using meteorological stations as described previously (Dussarrat *et al*. 2022). All metadata are available in Table S1.

### 2.3. Chemical extractions

To test method reproducibility, technical replicates were prepared for each rodent midden (Fig. 1). Middens used in this study were, in reality, the heart (*i.e.* the outer surface was removed in sterile conditions) of the previously discovered middens (Fig. S1). Under sterile conditions, each midden heart was bisected, and the hearts of these two subsamples were collected. Each tool used to divide the samples was washed with ethanol 70% between each step. Subsamples were ground individually twice for one minute using metal beads to obtain a thin powder (Fig. 1 and S1). To analyse primary metabolites, 8 mg of dried powder were extracted in 1 mL of 80% methanol (v:v) with 10 µM ribitol (99%, AppliChem) as an internal standard. The mixture was placed in a Precellys24 Instrument (VWR, Darmstadt, Germany), using 1 mm zirconia beads (Roth, Karlsruhe, Germany). Extracts were treated three times at 6.5 m/s for 45 sec. After 20 min centrifugation at 15,000 *g*, the 750 µl clear supernatant was transferred to 1,1 mL glass vials (Macherey-Nagel, Düren, Germany) and evaporated in a nitrogen stream. Metabolites were derivatised with 75 µL methoxylamine hydrochloride in pyridine (20 mg/mL; g/v) for 90 min at 37°C and 75 µL MSTFA for 30 min at 37°C. All chemicals and standard compounds were purchased from Roth (Karlsruhe, Germany), Sigma-Aldrich (Taufkirchen, Germany) or Macherey-Nagel (Düren, Germany).

**Fig. 1.**
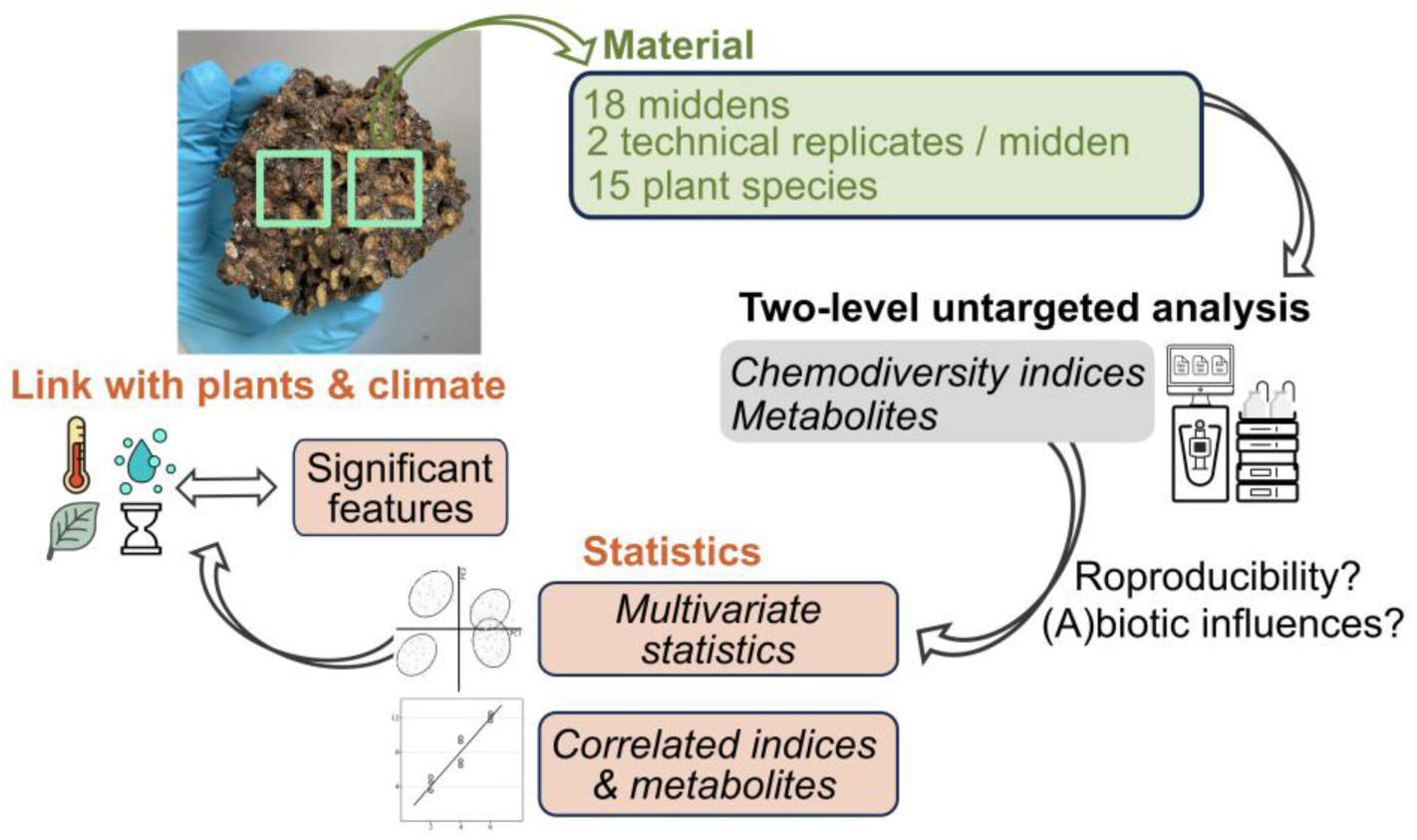
Simplified experimental framework.

For untargeted analysis, 10 mg of dried powder were extracted in 550 µL of a 90% methanol solution (v:v) with hydrocortisone as an internal standard (10 mg/L, Sigma-Aldrich, Steinheim, Germany). The mixture was homogenised on an automatic vortex for 5 min, sonicated in an ice bath for 15 min and centrifuged for 10 min. Supernatants were collected, filtered using 0.2 µm syringe filters (Phenomex, Torrance, CA, USA) and transferred into vials for LC-MS analysis (Dussarrat *et al*. 2023).

### 2.4. Metabolomics

A TSQ 9000 Triple Quadrupole GC-MS/MS (Thermo Electron, Dreieich, Germany) was used with an OPTIMA 5MS-0.25µM, 30m*0.25µM column (Macherey-Nagel, Germany). Samples were processed with an injection of 1 µl sample at 280°C with an ion source at 280°C, a helium flow of 1 mL/min, splitless mode, and a temperature gradient of 80°C for 3 min followed by an increase of 5°C/min to 320°C and a plateau for 2 min. The system was equilibrated for 2 min at 80°C after each analysis. Mass spectra were recorded at 1 scan/s with a scanning range of 50-750 m/z. Pre-processing was performed on Xcalibur (V.4.5, Thermo Electron, Dreieich, Germany). Identification was performed by comparing mass spectra and retention indices, which were defined according to an alkane standard mix (Sigma-Aldrich), to an in-house database. Peak areas of compounds with two analyte peaks were summed, and a normalisation by the peak area of the internal standard and sample weight was conducted to normalise the areas of the 67 compounds identified by comparison to purified standards. The injection of sample M6 failed and was therefore excluded from the table (Table S2).

To explore the chemical diversity, methanolic extracts were subjected to untargeted analysis via liquid chromatography (Vanquish Flex)-Orbitrap mass spectrometry (Exploris 120) with an electrospray ionisation source (ThermoScientific, Bremem, Germany) operating in negative mode. Separation was performed using a Kinetex XB-C18 column (150 x 2.1 mm, 1.7 µm, with guard column; Phenomenex) maintained at 45°C with a flow rate of 0.5 mL/min. Water (TH Geier – Chemsolute)-0.1% formic acid (FA) and acetronitrile-0.1% FA were used as solvents A and B with a previously established gradient (Dussarrat *et al*. 2023). MS/MS spectra were acquired at resolution 60,000 for the MS scan and resolution 15,000 for the MS/MS scan using data-dependent analysis with a range of 70-1000 m/z. The following MS parameters were used: lock mass correction EASY-IC, ion source type H-ESI, spray voltage 2500V, sheath gas 50 arb, aux gas 10 arb, sweep gas 1 arb, ion transfer tube temperature 325 °C, vaporiser temperature 350 °C, intensity threshold for MS/MS 5.0e^3^, HCD collision energy 30% normalised. To control the quality of the run and extraction process, quality control (QC, a pool of 12µL of each sample) samples were injected every 15 samples, and six extraction blanks were added.

Raw LC-MS data were processed via mzmine (v.4.1.0) using optimised parameters: minimum feature height of 9.10^4^, *m*/*z* tolerance of 10 ppm, *m*/*z* tolerance intra and inter-samples of 3 and 5 ppm, Savitzky Golay as smoothing algorithm and a mass detector set up as Factor of lowest signal, retention time tolerance intra- and inter-sample of 0.04 and 0.1 min (Schmid *et al*. 2023). Subsequently, peak intensities were divided by the internal standard and features were removed if: coefficient variation in QC > 30% or intensity 10 times lower than average extraction blank intensity, yielding 9,550 features. Peak intensities were then related to sample weight (Table S3). LC-MS and GC-MS datasets were normalised using median normalisation, cube-root transformation and Pareto scaling on MetaboAnalyst v.6 (Pang *et al*. 2024) prior to statistical analysis as previously described (Dussarrat *et al*. 2022, 2025) (Tables S4 and S5, and deposited online (see Data availability). Putative chemical formulas, chemical families (which include Natural product classification (NPC) pathways, superclasses and classes) and putative compound names were defined using SIRIUS (v. 6.1.0) with Orbitrap default parameters (5 ppm, de novo plus bottom up strategy below 500 Da) (Dührkop *et al*. 2019, 2021; Hoffmann *et al*. 2022). The most probable pathway, superclass and class of each feature were assigned only if the confidence score was ≥ 0.8, as recommended (Hoffmann *et al*. 2022). Confidence in the annotation level for each chemical feature was assigned according to the Metabolomics Standards Initiative (Sumner *et al*. 2007). Metabolic networks were developed by including the mzmine output files into GNPS using the feature-based molecular networking pipeline with a mass tolerance of 0.02 Da, a minimum paired *cos* score of 0.7 and a minimum matched fragment ions of 5 (Nothias *et al*. 2020; Wang *et al*. 2016). Networks were then customised using Cytoscape (v.3.10.4) (Shannon *et al*. 2003).

To calculate chemodiversity indices, Shannon diversity and Functional Hill (FH) diversity were defined using the *chemodiv* package on R (v.4.5.1) (Petrén *et al*. 2023; R Core Team 2024). The cos matrix was extracted from GNPS to develop the dissimilarity matrix (1-cos score) required to define FH diversity for each chemical family. Chemical families that included less than 0.1% of the total pre-processed features (< 10 features; 0.1%*9550) were excluded to simplify the interpretation and limit potential false positives. The resulting table included 300 chemical indices, which were normalised (median normalisation, log 10 transformation and autoscaling) prior to data analysis (Brückner & Heethoff 2017; Petrén *et al*. 2023) (Tables S6 and S7).

### 2.5. Statistical analyses

To test method reproducibility, we explored Bray-Curtis distances and Chemical Structural and Compositional Similarity (CSCS) distances between pairs of midden samples and between technical replicates (Bray & Curtis 1957; Sedio *et al*. 2017). Bray-Curtis distances were calculated using the *vegdist* function from the *vegan* package (Oksanen *et al*. 2025). CSCS distances were estimated by submitting the similarity matrix to the *cscs* function of the *rCSCS* package, as previously described (Brejnrod *et al*. 2019; Nomoto *et al*. 2025; Sedio *et al*. 2017). To estimate the influence of metadata parameters, Bray-Curtis and CSCS distance matrices were then visualised on NMDS plots created using the *vegan* package, where the significance of the effect of metadata parameters was tested using the *envfit* function (Oksanen *et al*. 2025).

To explore the influence of midden composition (*i.e.* number of species, phylogeny distance), age and abiotic parameters (*i.e.* temperature, CO_2_, precipitation), we separated plant and midden samples. Correlations between metadata parameters and chemical feature intensities or chemical indices were primarily calculated using midden samples. First, we calculated the average intensity of each chemical feature of chemical indices between midden technical replicates (Tables S2, S5, and S7). Next, correlations were calculated using *Hmisc* package and the *p.adjust* function (Jr 2025). Importantly, chemical indices and features were considered as significantly influenced by age, temperature, CO_2_ or precipitation only if *P* < 0.05 and no significant association with midden composition (number of species or Faith’s index). To illustrate the results, heatmaps, boxplots and barplots were designed with *pheatmap*, *cluster*, *factoextra* and *ggplot2* packages (Kassambara & Mundt 2020; Kolde 2025; Wickham 2016) and Venn diagrams were designed on https://bioinformatics.psb.ugent.be/webtools/Venn/. Next, we used Partial Least Squares regression (PLSr) to test the capacity of midden chemistry to fit midden age, CO_2_ and temperature and precipitation anomalies. Models were run using the *pls* package on R with the leave-one-out validation method and two components (Dussarrat *et al*. 2025; Liland *et al*. 2024). We then tested whether chemical features and chemical indices responding significantly to temperature, CO_2_ and precipitation in midden samples were also influenced by temperature, soil water content and elevation in plants (Fig. 1). Finally, to test whether midden chemistry could be used to track species composition of the middens, we performed Orthogonal Partial Least Squares-Discriminant Analysis (OPLS-DA) using MetaboAnalyst (v.6) (Pang *et al*. 2024). R scripts used in this study were deposited online (see Data Availability section).

## 3. Results 1200-1500 words

### 3.1. Excellent capacity of untargeted metabolome to explore the metabolism of ancient midden samples

As a first step to explore the possibility of conducting untargeted metabolomics on ancient midden samples, we compared plant and midden chemistry. Consistent with their mixed biological origin, middens captured 79% of the chemical diversity of all 15 plant species used in this study (Fig. 2A). Moreover, 65% (5,574) of detected features were found in both biological matrices, while 29% and 17% were exclusive to middens and plants, respectively. In addition, chemical richness (*i.e.* number of detected features) showed greater variation between middens than between plant samples (Fig. S2). Next, we tested the reproducibility of the approach. Bray-Curtis dissimilarities between middens were considerably higher (ratio of 9 between medians) than between technical replicates, supporting the reproducibility of the method (Fig. 2B).

**Fig. 2.**
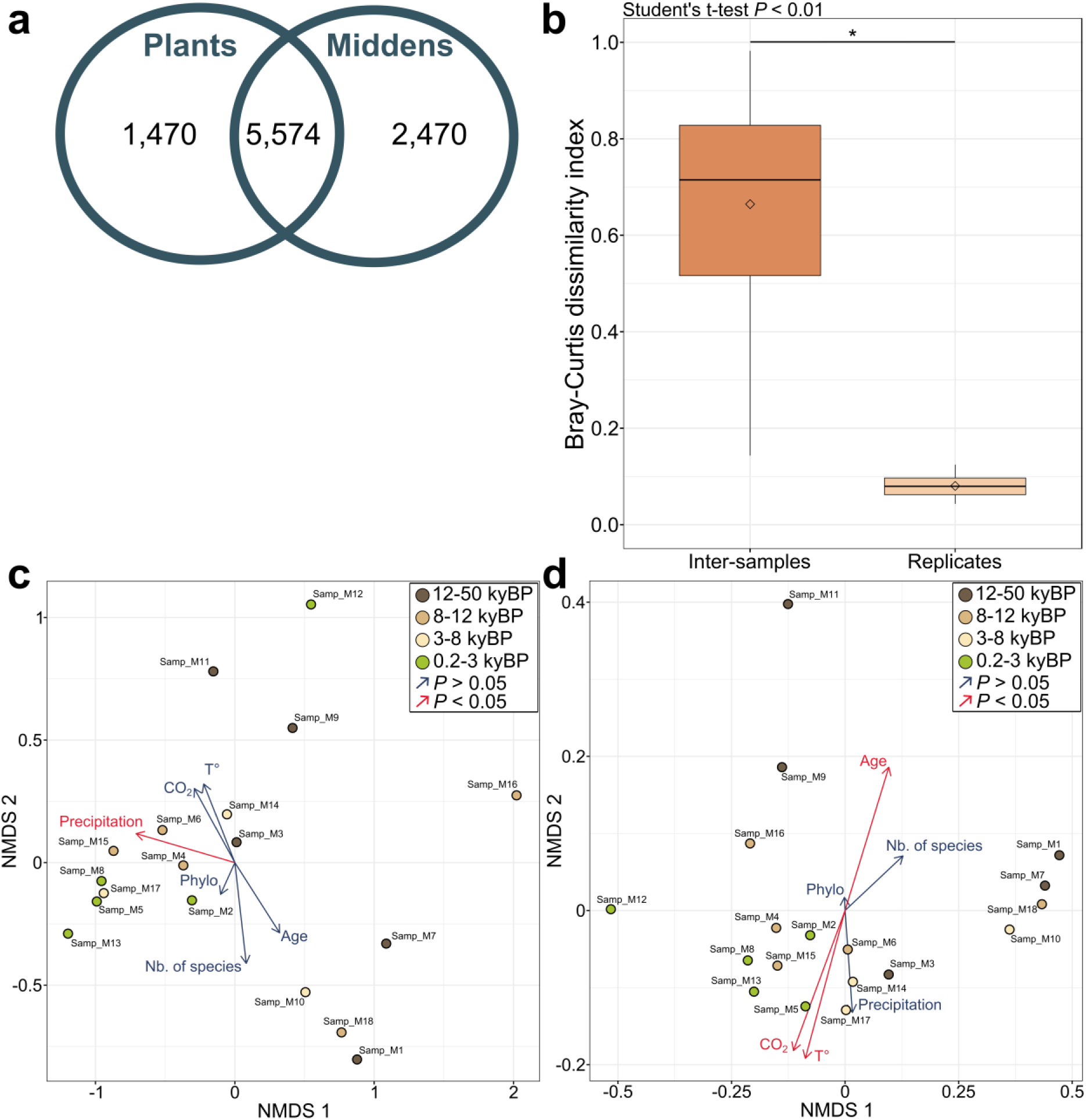
Chemical distances between samples. **A.** The number of detected features represents the number of pre-processed ions that were detected in at least one midden or one plant. **B.** Significant variation in Bray-Curtis dissimilarities between distinct samples or between technical replicates (Student’s t-test, *P* < 0.01). **C-D.** Non-metric multidimensional scaling analysis (NMDS) using (**C**) Bray-Curtis distances or (**D**) chemical structural compositional similarities (CSCS). Red arrows represent environmental factors that have a significant impact on the Bray-Curtis (none were significant) or CSCS distances (*P* < 0.05). Blue arrows refer to non-significant effects. For each NMDS, the influence of temperature, precipitation, CO_2_, age, number of species in the middens, and Faith’s phylogenetic index was tested. Samples were coloured according to age. *kyBP: kiloyear before present*.

### 3.2. Distinct consequences of midden composition, ageing and climate on midden chemistry

The variation in total richness observed in midden samples was not correlated with midden plant composition (*i.e.* number or phylogenetic diversity of plant species included in the midden) nor with age (Fig. S2). Although age did not have a significant impact on total richness, it did have a notable influence on Bray-Curtis and CSCS distances (Fig. 2C and 2D) and, consequently, on midden chemistry. This result was corroborated by PLS-DA, which showed distinct patterns between the different age groups and by a heatmap of midden chemistry at the chemical feature level (Fig. S3 and S4). Although the distribution of midden samples differed slightly depending on whether Bray-Curtis or CSCS distances were used, both approaches suggested a distinct influence of (i) age, (ii) precipitation, and (iii) temperature with CO_2_ on midden chemistry (Fig. 2C and 2D).

### 3.3. Richness, diversity and disparity of major chemical families fluctuate with climate

Next, we characterised the influence of midden plant composition, age and climatic factors on the chemodiversity of midden samples (Fig. 1). Midden plant composition had a limited effect on chemodiversity, significantly influencing (*P* < 0.05) the disparity and diversity of coumarins, oligopeptides and apocarotenoids, for instance (Table S8). To minimise false positives, these significant indices were excluded (but still available in Table S8) when examining the effects of age and climate. A total of 69 chemical families (28 with FDR correction) were influenced by age, CO_2_, precipitation, and temperature, respectively (Fig. 3A). Overall, the three components of chemodiversity responded similarly (Fig. S5). The combined effect of temperature-CO_2_ was again highlighted (Fig. 3B), where a decrease in temperature was followed by an increase in diversity of flavonoids, lipids (*e.g.* eicosanoids), a decrease in polyketide diversity, and an increase in phenylpropanoid richness, for instance (Fig. 3C and Table S8). The midden samples available for this study were produced during very wet periods (compared to current days), as indicated by strongly positive precipitation anomalies (Fig. 3C and Tab. S1). High to very high precipitation levels were associated with higher diversity of sugars and several phenolics (*e.g.* lignans, flavones), and higher richness of megastimanes.

**Fig. 3.**
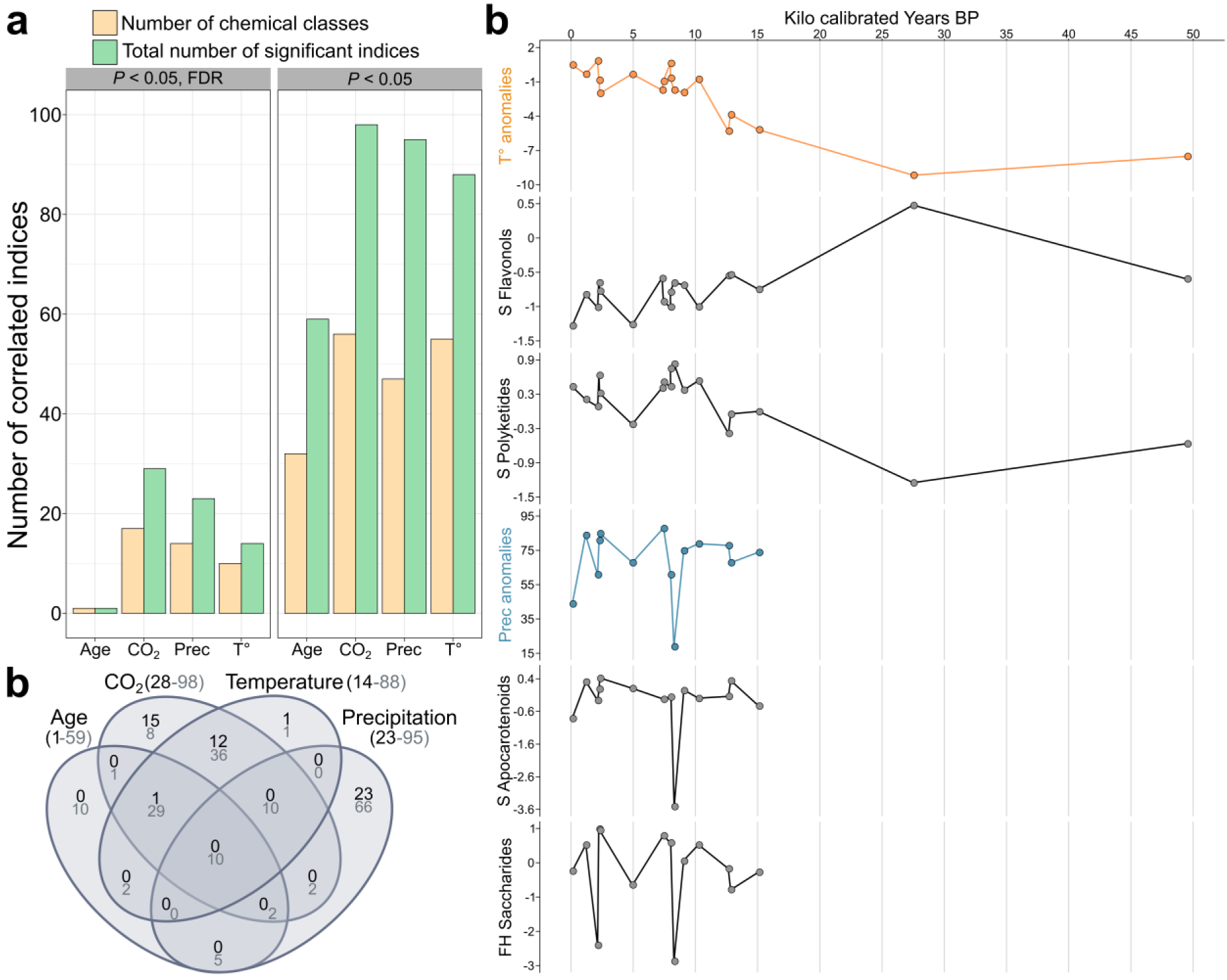
Influence of temperature and precipitation anomalies and CO_2_ on chemical indices. **A.** Depiction of the significant chemical indices. *Prec*: *precipitation anomalies; T°*: *Temperature anomalies.* **B.** Venn diagram of the chemical indices showing significant variations in response to age, temperature, CO_2_ or mean annual rainfall. Chemical indices that were also significantly (*P* < 0.05, no FDR) linked to either the number of species in the midden or to Faith’s diversity index of the midden were excluded from the figure. Dark numbers: correlation *P* < 0.05, FDR; grey numbers: correlation *P* < 0.05, no FDR correction. **C.** Examples of chemical indices significantly linked to temperature or precipitation anomalies. *T°: temperature; Prec: precipitation; R: richness; S: Shannon diversity; InvSi: inverse Simpson index; FH: Functional Hill diversity*.

### 3.4. Chemical response at the metabolite level can trace midden composition, age and climate

To gain further insights into how midden chemistry reflected a changing environment, we explored variation at the metabolite level (Fig. 1). The study of primary metabolites highlighted an effect of midden composition on hydroxyproline, malate, fumarate, and homoserine, for instance (Table S9). Lower temperatures were associated with higher levels of aspartate, pyroglutamate, glycine, β-alanine and 2-hydroxyglutarate and lower levels of adenosine and asparagine. In contrast, very high precipitation levels observed during this period were linked to higher contents of glycerol-3-phosphate, sucrose and aspartate (Fig. S6 and Table S9). The influence of age and climatic parameters on the richness and diversity of major chemical families was also reflected in the analysis of the metabolic fingerprints (Fig. 4 and Table S10). Overall, 135 features were influenced by midden composition and 1,823 features by climatic parameters or age (excluding features related to midden composition). In line with results at the chemodiversity level, most significantly affected features (*P* < 0.05, FDR) responded to temperature and CO_2_ on one side and precipitation on the other (Fig. 4A). Besides, several clusters of chemical features, which included terpenoids, fatty acids and phenolics, also showed substantial variation in response to these abiotic factors (Fig. S7 and S8). The interest in these pathways was reinforced after the annotation of a set of features composed of the top hundred correlated features to age, temperature, CO_2_ and precipitation levels (Table S11, which also included features observed in relevant clusters (Fig. S7 and S8)). The pathways “Shikimates and phenylpropanoids” and “Fatty acids” accounted for 32% and 23% of this set of annotated chemical features, supporting their role in response to temperature and precipitation pressures, as previously shown in the study of chemical indices (Fig. 4B). Fatty acids were mostly influenced by temperature and precipitation (36 and 62% of annotated significant fatty acids, respectively), while phenolics were mostly influenced by precipitation levels (70%). Although less observed in the study of chemodiversity, terpenoids (*e.g.* diterpenes and sesquiterpenes) showed great variation in response to age, temperature and precipitation.

**Fig. 4.**
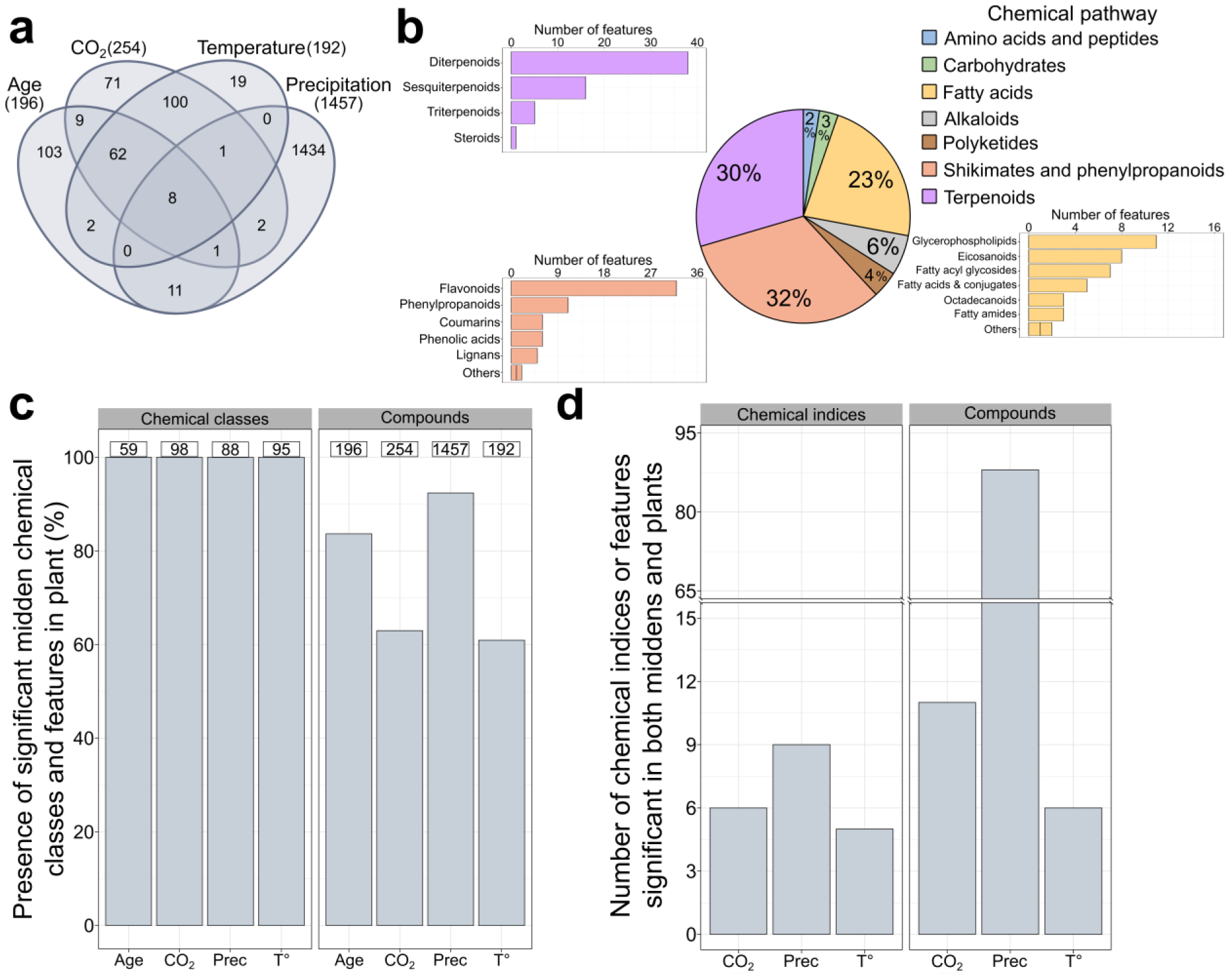
Performance of midden metabolism to integrate environmental constraints and supporting results on modern plant samples. **A.** Venn diagram of the chemical features showing significant variation in function of age, temperature, CO_2_ or precipitation (*P* < 0.05, FDR). **B.** Annotation and classification of the most discriminant features. Chemical features included in this graph were: the top 100 features correlated with temperature and precipitation anomalies, age and CO_2_, as well as features significantly correlated to abiotic parameters in both plant and midden samples (see Table S12 for the complete annotation table). **C.** Presence of significant chemical classes (*e.g.* Saccharides) and compounds in plants. Plants were collected across an elevation gradient from a similar location as the middens (see Methods section). *Prec: precipitation anomalies; T°: temperature anomalies*. **D.** Number of chemical indices or features significantly correlated in both middens and plants. *CO_2_: chemicals correlated with CO_2_ in middens and elevation for plants. Prec: chemicals correlated with precipitation anomalies in middens and soil water content in plants. T°: chemicals correlated with temperature anomalies in middens and temperature in plants*.

We then tested the capacity of midden chemistry to trace age and climatic variation. Midden chemistry fit the midden age with an *R²* of 0.9 (Fig. S9). As expected with the detection of significant chemical features, temperature, CO_2_ levels and precipitation anomalies also fit with a minimal *R²* of 0.88 (Fig. S9). Besides, midden chemistry could be used to survey the presence of plant species over time, successfully distinguishing middens that contain or do not contain specific plant species (Fig. S10).

### 3.5. Significant indices and chemical features in middens are congruent with the chemical response of recent plants to elevation

To provide insights into whether the uncovered chemical response more likely originated from plant material or other potential biological sources, we explored the plant response to elevation, temperature and soil water content across an elevation gradient. All chemical families that included significant indices in middens were observed in plant samples, as well as 88% of the significantly affected chemical features (Figure 4C and Table S12). Although the majority of the remaining 12% of features were linked to temperature and/or CO_2_, 60% of all chemical features related to these abiotic pressures were found in plant samples. In total, 20 chemical indices and 105 chemical features were significantly correlated to variation in abiotic parameters in both plant and midden samples (Fig. 4D). Soil water content and temperature influenced the chemodiversity of multiple chemical families in sugar, lipid and phenolic pathways, which were also influenced by precipitation levels and temperature in midden samples (Fig. 4D and Table S12). Similar observations were made at the compound level, where water availability and temperature affected primary metabolites, particularly fatty acids (*e.g.* glycerophospholipids), and specialised features, particularly phenolics (*e.g.* flavonoids, lignans) and some terpenes. Overall, the response of lipids was consistent (*i.e.* positive correlation with precipitation and soil water content) across both biological matrices, while opposite for specialised chemical families (Table S12).

## 4. Discussion

Through millions of years of evolutionary pressure, every organism has forged the metabolism best suited to thrive in its own environment (Scossa & Fernie 2020). Besides, given the species specificity of metabolism and its responsiveness to environmental variation, exploring ancient metabolism could offer exciting opportunities to reconstruct past ecosystems and temperature or precipitation patterns (Badillo-Sanchez *et al*. 2023; Dussarrat *et al*. 2022; Scossa & Fernie 2020). Despite this potential, the use of paleometabolomics is remarkably scarce, and applied almost exclusively to studying human-related questions (Cole *et al*. 2025). Here, we tested and validated the possibility of performing non-targeted metabolomic analyses on ancient samples in a reproducible process. Using a timeline of rodent middens across the last ca 50 thousand years, we highlighted major changes in chemodiversity of various chemical families and showed the potential of midden metabolism to predict ecosystem composition and changes in temperature, CO_2_ and precipitation patterns. Shifts in chemical families and metabolites in midden samples were then tested on plants collected along an elevation gradient at the midden sites.

### 4.1. High efficiency and reproducibility of untargeted metabolomics on ancient samples and limitations

A major challenge for untargeted metabolomics is the need to capture a representative fraction of the sample. Rodent middens can vary in size but can reach tens of cm in diameter, questioning the possibility of achieving reproducible grinding and homogenisation (Becklin *et al*. 2024; Latorre *et al*. 2003). Our results showed very limited chemical dissimilarities between technical replicates, which were obtained by taking two independent subsamples (cores) from each midden, supporting the reproducibility of our workflow (Fig. 2). Besides, midden metabolism captured 79% of the chemical diversity detected in 15 plant species recently collected near midden samples, showing the high efficiency of untargeted metabolomics on ancient samples (Zimmermann *et al*. 2021) and supporting the midden capacity to capture the diversity of the surrounding flora (Becklin *et al*. 2024). Overall, middens have emerged as an ideal material for combining metabolomics and paleoecology to infer past ecosystem dynamics and climates, offering a complementary strategy to herbarium studies. Herbaria offer rare opportunities to explore past metabolic survival strategies, but the number of collections is limited, and some often lack the geographical and historical context needed to link those shifts to ecological function (Eckert *et al*. 2025). Rodent middens can complement these disadvantages, but their analysis also carries some limitations. Rodent middens are composite deposits dominated by fecal pellets composed largely of plant remains, cemented within a matrix of crystallised urine that can entrap additional biological material (e.g. pollen, plant fragments, arthropod remains, microbial remains, dust), making it difficult to parse plant, insect or microbial defence strategies (Fig. S1). Together with a previous study (Díaz *et al*. 2019), our results suggest that exogenous contaminants can be largely excluded by working under sterile conditions and sampling the midden core. Furthermore, using recent samples to corroborate observations from ancient material is valuable for assessing whether the uncovered past metabolic strategies remain viable in contemporary organisms (Jackson & Williams 2004). Here, shifts in midden chemistry due to temperature and water availability variation were also tested on plants, highlighting chemical families and features that responded in both matrices (Fig. 4). Although a consistent correlation between climate and both midden and plant chemicals supports the role of these chemicals, a lack of validation (i.e. either an opposite pattern or absence of correlation) in recent plants does not imply a false positive, as ancient and recent plants may operate under different metabolic regulations (Jackson & Williams 2004). While plants represent valuable material for supporting our findings in ancient middens, gathering rodent middens of similar ages along the altitudinal cline would provide an ideal but extremely time-consuming validation system (Becklin *et al*. 2024). Altogether, our results support the efficiency and reproducibility of the developed workflow for untargeted metabolomics on ancient samples.

### 4.2. Midden chemistry is influenced by its intrinsic properties and climatic variations

Results from the different analyses converged to reveal the impacts of midden intrinsic properties and climate on the primary and specialised metabolites of rodent middens. First, the distinct effects of age, temperature-CO_2_ and precipitation levels were shown through Bray-Curtis and CSCS distances (Fig. 2). Although both analyses displayed similar outcomes, significant factors changed, emphasising the need to use both distances, as CSCS incorporates structural similarities but only considers the fraction of the chemical composition with MSMS data (Nomoto *et al*. 2025; Sedio *et al*. 2017). Richness, diversity and disparity of multiple chemical classes fluctuated with temperature and precipitation levels. In middens’ primary metabolism, decreased temperatures were associated with an increase in lipid diversity (*e.g.* eicosanoids), while lipid richness (*e.g.* glycerophospholipids) decreased with precipitation. The relevance of eicosanoids and glycerophospholipids was confirmed at the compound level. In addition, their role was further supported by increased contents under lower soil water content in plants. This link between lipids and hydroclimate variation is congruent with previous analyses of leaf waxes in rodent middens (Frugone-Álvarez *et al*. 2023). These findings broaden the observation to other lipid families, expanding the lipid repertoire that can be used to trace past changes.

In addition, results from GC-MS analysis revealed changes in several stress-related metabolites, such as amino acids, which were linked to specialised metabolism, and pyroglutamate (also called 5-oxoproline) (Singh *et al*. 2025; Trovato *et al*. 2021). Next, the shift in primary metabolism was followed by consequent changes in specialised chemical families. In midden samples, lower temperatures were associated with increased diversity of phenolics (*e.g.* flavonoids, phenylpropanoids), while higher precipitations were linked to higher diversity of lignans, flavonoids and megastigmenes. Flavonoids and phenylpropanoids typically accumulate under flooding (Li *et al*. 2021; Wang *et al*. 2019) and freezing (Díaz *et al*. 2024; Schulz *et al*. 2015) conditions, likely due to their defensive roles in modulating reactive oxygen species homeostasis (Agati *et al*. 2012), for instance. Lignans may also provide antioxidant protection and play a role in fortifying cell walls to face biotic threats, which may increase with higher precipitation levels (Šamec *et al*. 2021). In contrast, megastigmenes are degradation products of carotenoids (Samra *et al*. 2024), a process known for its link with phytohormones and redox signalling (Havaux 2014).

Interestingly, while lipid intensities rose with precipitation in both middens and recent plants, specialised metabolites (*e.g.* phenolics) tracked temperature consistently but showed opposite precipitation responses between the two (Table S12). For example, lower precipitations were associated with higher phenolic contents and diversity in plants. These patterns reflect contrasting stresses, as ancient plants in midden samples probably faced flooding during the Central Andean Pluvial Event (CAPE) I and CAPE II periods, while current plants are facing drought in the Atacama Desert (De Porras *et al*. 2017). Overall, our results support the central place of specialised metabolite diversification and regulation (at the compound intensity level) as an important and dynamic niche dimension (Müller & Junker 2022).

### 4.3. Results support potential applications of paleometabolomics on middens

Our results highlight paleometabolomics as a powerful tool with various applications from ecology to agriculture and services to humankind. First, we showed the excellent capacity of metabolism to track climate changes. The overdispersion or increased intensities of defensive compounds, such as redox-related compounds (*e.g.* phenolics), could be used to infer climate variation over the last 50,000 years. Once combined with predictive metabolomics, this approach could be used to support and even refine paleoclimate models (Dussarrat *et al*. 2022). Besides, depicting past changes in the metabolism of living species would improve our understanding and predictive capacity of current metabolic adaptations to climate change (Fordham *et al*. 2020). These actions would be beneficial at different scales: providing innovative strategies for crop engineering, serving biodiversity protection, and protecting biological resources for drug discovery (Castañeda-Álvarez *et al*. 2016; Defossez *et al*. 2021; Fordham *et al*. 2020). Moreover, while variation in chemical families and compound intensities was associated with midden composition or climate fluctuations, midden chemistry fitted midden age with an *R²* of 0.9. This chemical shift in response to age has been previously observed (Velsko *et al*. 2017). Hence, these observations raise the question of whether paleometabolomics could complement ^14^C dating as an independent temporal signal, while also providing a molecular alternative when DNA preservation is limited after the Middle Pleistocene (Brunson & Reich 2019; Dalén *et al*. 2023). However, additional studies are needed as the persistence of chemicals into deep time is yet to be explored. Finally, middens’ chemistry predicted the presence and absence of plant species. Leveraging the specificity down to the species scale of specialised metabolites, paleometabolomics could help reconstruct past ecosystems, from animals and plants to microbial communities. Such an approach could be used in complement to pollen assemblage analyses, which are limited by the seasonal and species-specific nature of pollen production, or coprolite analyses via paleogenomics (Marquer *et al*. 2014; Wood *et al*. 2013).

## 5. Concluding remarks

Paleometabolomics offers exciting prospects for exploring life’s ancient metabolic innovations to meet climate change, yet the potential for untargeted metabolomics on ancient samples remains to be defined. Here, we used a discontinuous timeline of rodent middens up to 49,600 years old to test the capacity of untargeted metabolomics to reproducibly capture the chemistry of ancient samples and investigate the impact of climatic variations. Our results support the use of paleometabolomics, as middens’ chemistry could be described in a reproducible way and captured 79% of the chemical richness of 15 plant species collected recently near the midden sites (Dussarrat *et al*. 2022; Latorre *et al*. 2002). Exploring variation in midden chemistry could effectively fit climate variation and plant species presence, supporting the use of paleometabolomics to reconstruct past ecosystems and explore how life navigated past climate events (Badillo-Sanchez *et al*. 2023; Becklin *et al*. 2024). Lipids and phenolics (*e.g.* lignans, flavonoids), which responded significantly in both midden and plant samples, emerged as compounds of interest for tracking temperature and precipitation dynamics in ancient samples.

## Author contributions

TD, CL and RG designed the experiment. TD, FD, CC, CR, PP, CL and RG performed the sampling. TD, VG, MP, and KN performed chemical extractions and GC-MS and LC-MS analyses. TD analysed the data, and TD, FP, VG, MP, PP, CM, CL and RA interpreted the results. TD wrote the first draft of the manuscript with the feedback of all co-authors.

## Acknowledgements

CM and TD acknowledge the financial support from the German Research Foundation (DFG, project MU 1829/28-2). TD thanks the Faculty of Biology of Bielefeld University for financial support (Project I-3210-0213-0014, Forschung und wissenschaftlicher Nachwuchs). We are grateful to the indigenous community of the Comunidad Indígena de Talabre for the access to the transect. RG and CL acknowledge funding from the ANID Millennium Scientific Initiative Program (iBio ICN17_022; CGR ICN2021_044), and IEB (FB210006). FPD acknowledges AFOREST (NCS2022_24).

## Data Availability Statement

Data will be made available upon publication of the study.

