## Supplemental Figures for "Paleometabolomics reveals impacts of abiotic factors on rodent midden metabolism over the last 50,000 years"

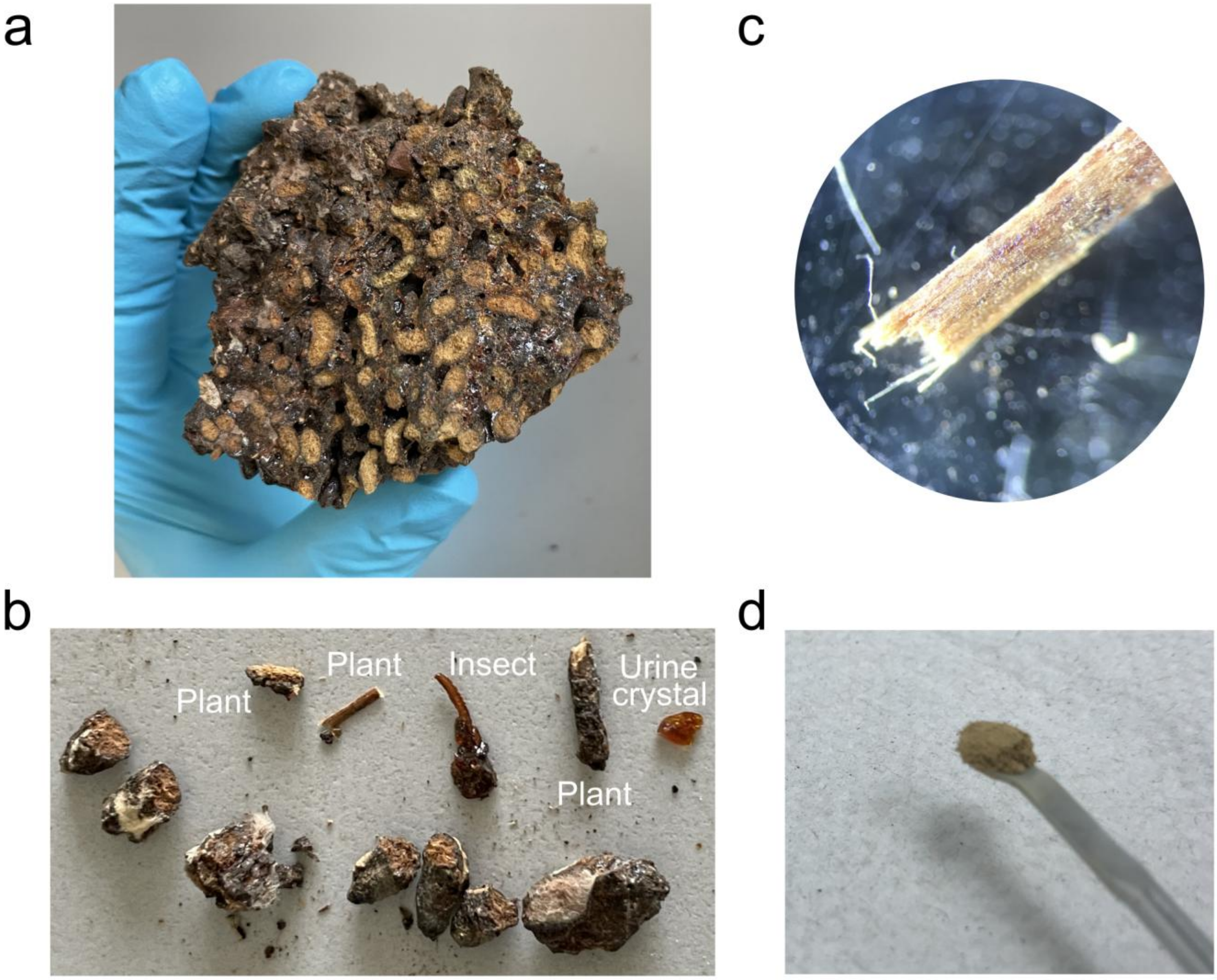


**Fig. S1 | Rodent midden composition. A**. Rodent midden. B. Different parts included in the midden, such as plant material, urine crystals, and insect legs. **C.** Binocular view of plant stems in a rodent midden dating back more than 10,000 years. **D.** Midden powder obtained after grinding.

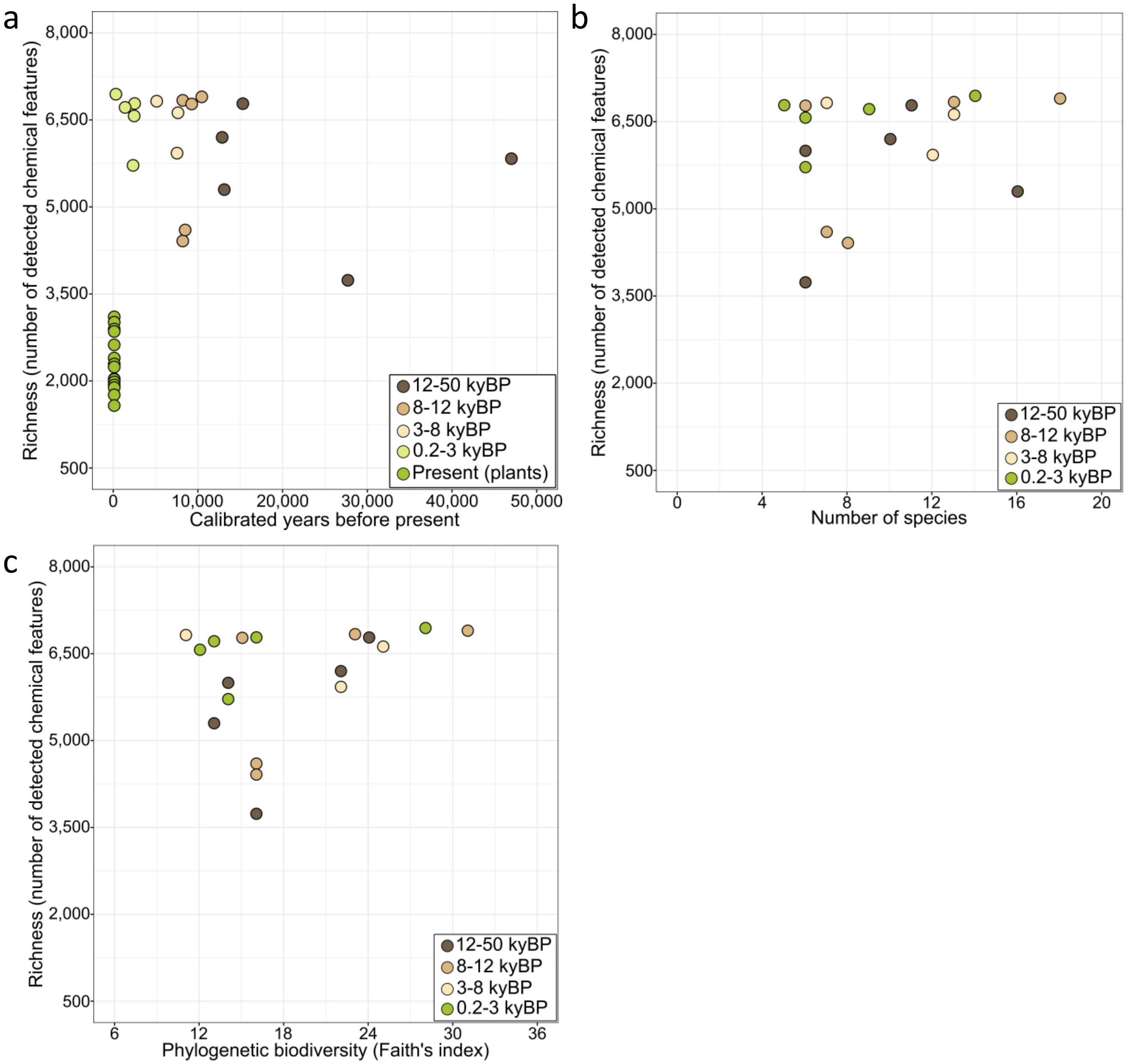


**Fig. S2 | Absence of links between chemical richness and age or midden biological composition. A-C**. Depiction of the links between feature richness (i.e., the number of chemical features detected) and (**A**) age, (**B**) the number of species present in middens, and (**C**) midden phylogenetic biodiversity.

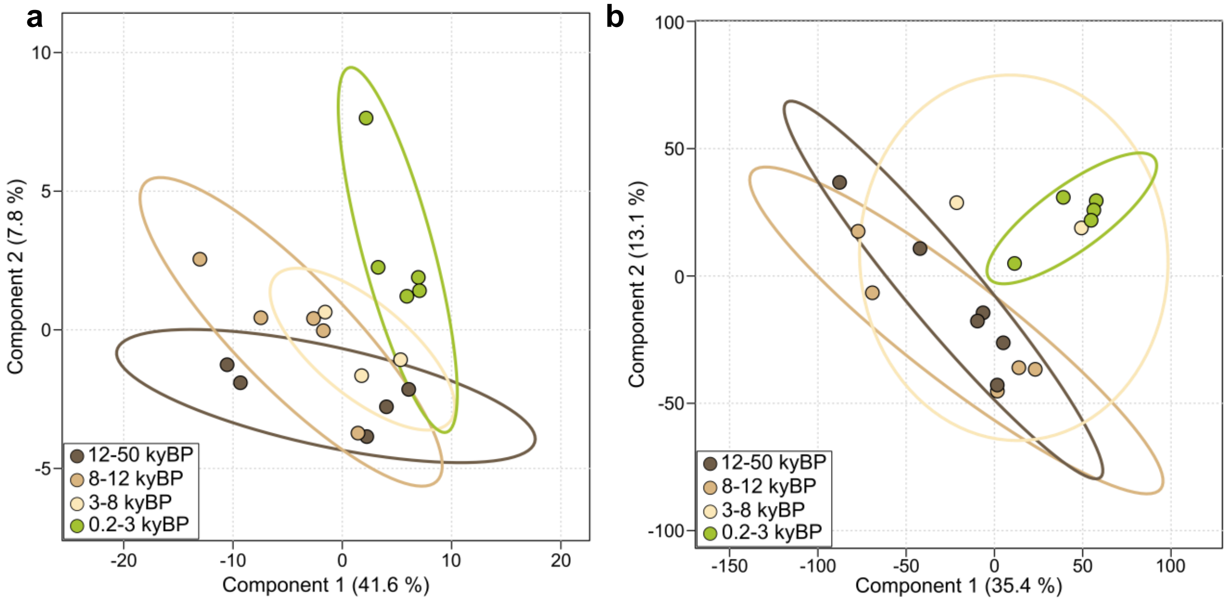


**Fig. S3 | Variation in middens' chemistry. A-B.** Partial least square discriminant analysis (PLS-DA) using (**A**) chemical indices (300) or (**B**) chemical features (9,514). *kyBP: kiloyear before present.*

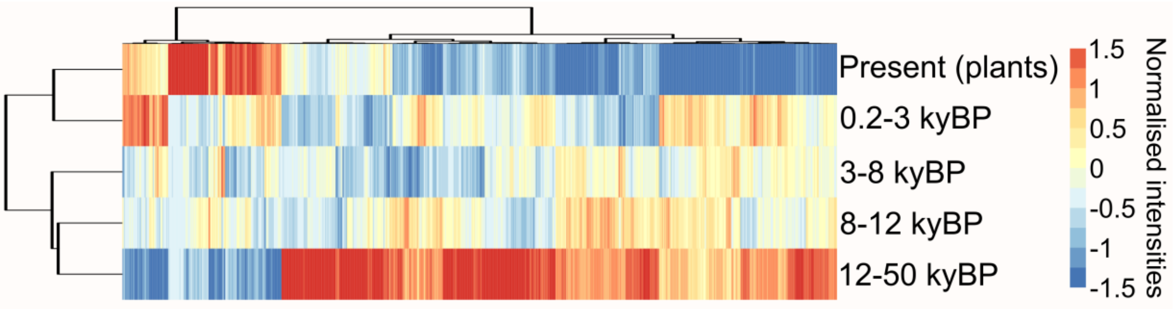


**Fig. S4 | Chemical variation in midden samples at the metabolite level.** Heatmap of the chemical composition of the samples (9,514 chemical features).

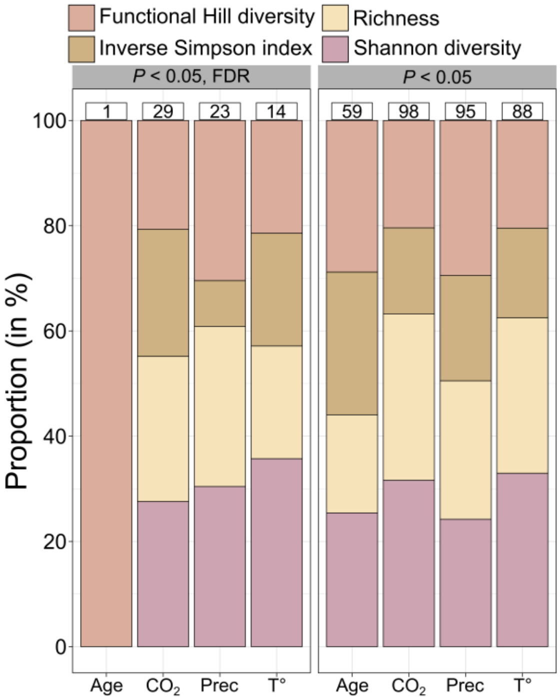


**Fig. S5 | Depiction of the significant chemical indices.** Depiction of the type of chemical indices responding significantly ($P < 0.05$, with or without FDR correction) to the different parameters. Numbers in the square refer to the total number of correlated indices. *Prec*: *precipitation anomalies; T°*: *Temperature anomalies.*

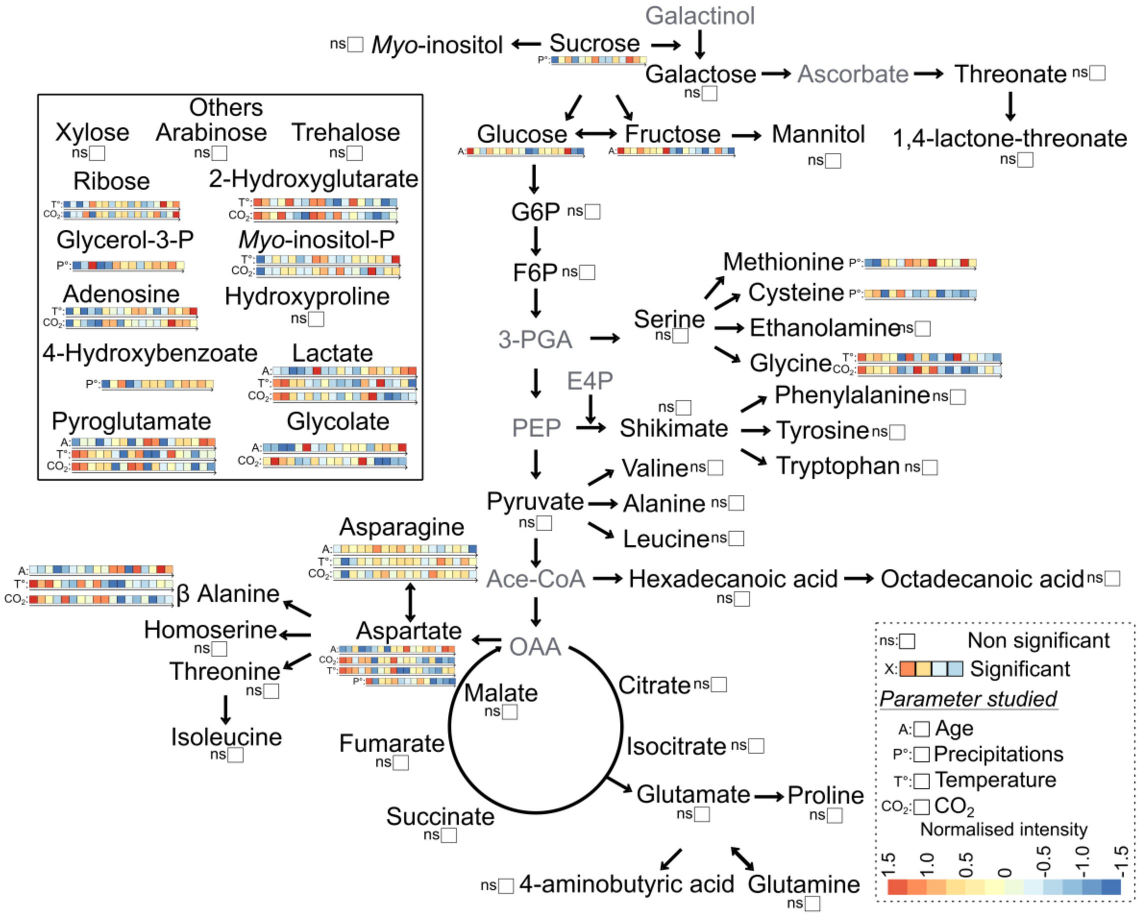


**Fig. S6 | Influence of age and abiotic parameters on primary compounds.** Simplified overview of the detected compounds via GC-MS. Grey compounds were not detected in our analysis. Heatmaps were linked to the compounds that showed significant variation in function of either age, temperature anomalies, $CO_2$ or precipitation anomalies. Compounds that were also significantly associated with the phylogenetic index (*i.e.* Faith's index) or the number of species detected in the midden were noted as non-significant. For more information, see Table S8.

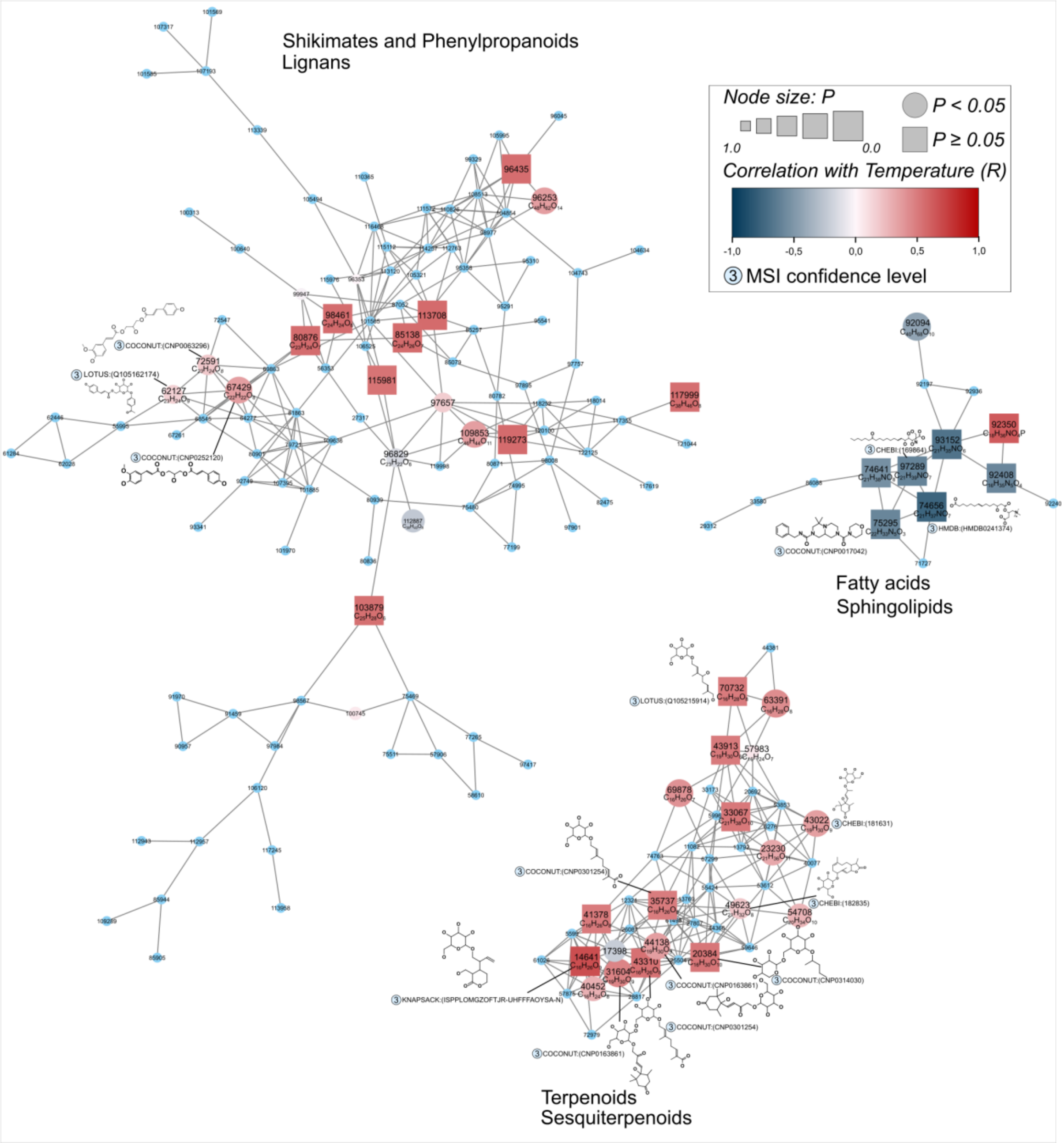


**Fig. S7a | Chemical clusters responding to temperature.** Example of clusters of chemical features responding to temperature anomalies. For more information, see Tables S10 and S12. Blue ellipses represent ions that did not successfully pass preprocessing. Chemical structures were drawn using Pubchem. The top 1 match from SIRIUS was used to draw the figure (MSI 2 or 3). Chemical pathways and superclasses represent the most represented ones in a given cluster. In this figure, annotations are putative and should be interpreted with caution at the compound level.

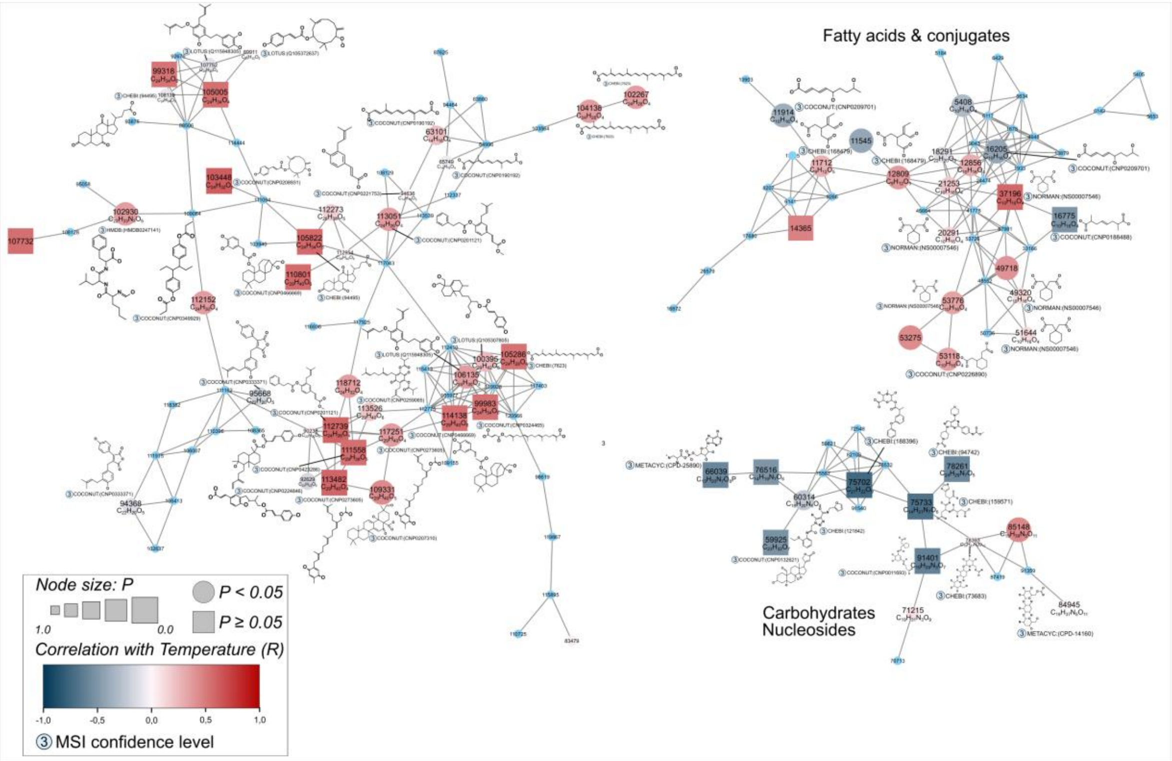


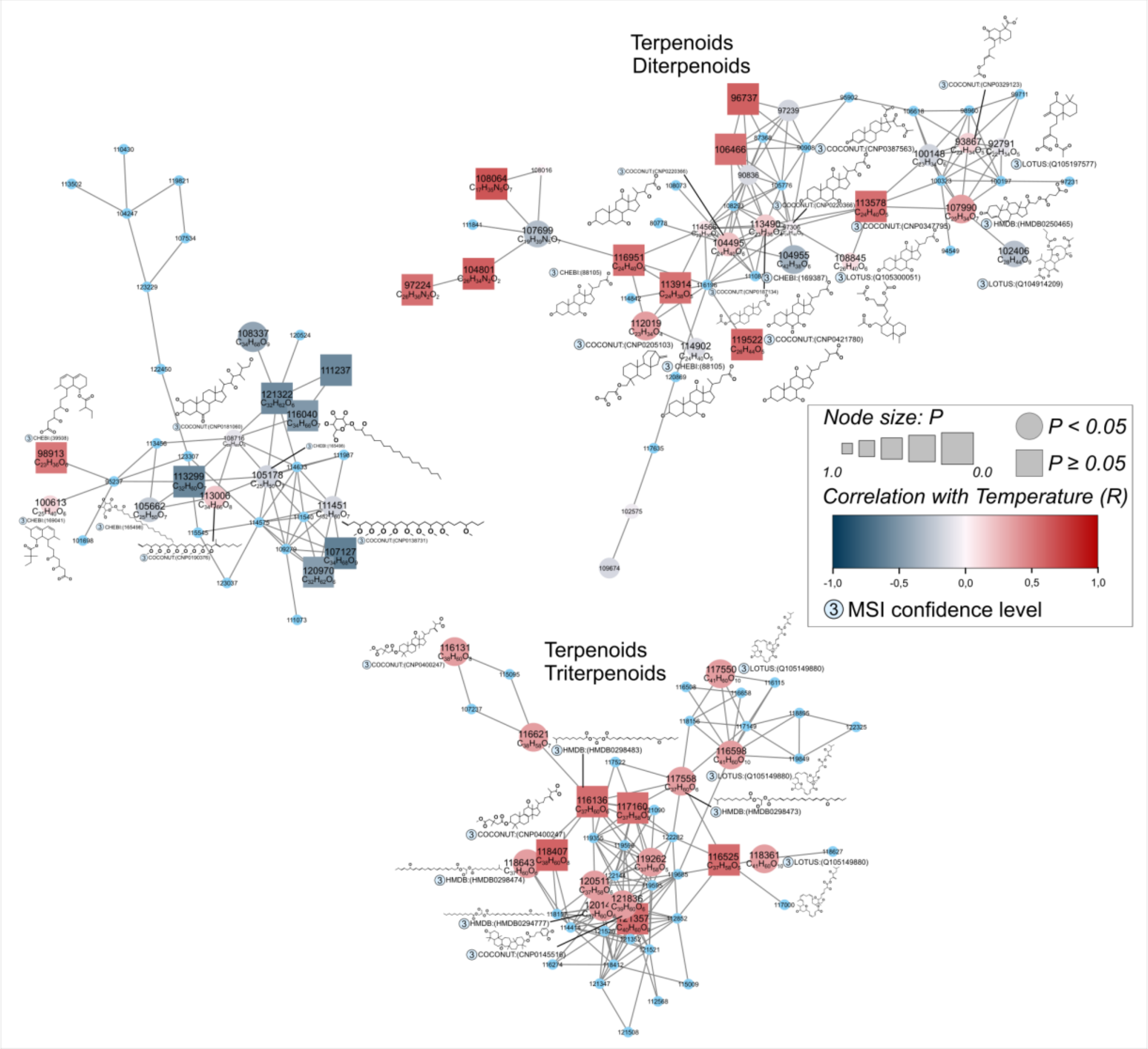


**Fig. S7b | Chemical clusters responding to temperature.** Example of clusters of chemical features responding to temperature anomalies. For more information, see Tables S10 and S12. Blue ellipses represent ions that did not successfully pass preprocessing. Chemical structures were drawn using Pubchem. The top 1 match from SIRIUS was used to draw the figure (MSI 2 or 3). Chemical pathways and superclasses represent the most represented ones in a given cluster. In this figure, annotations are putative and should be interpreted with caution at the compound level.

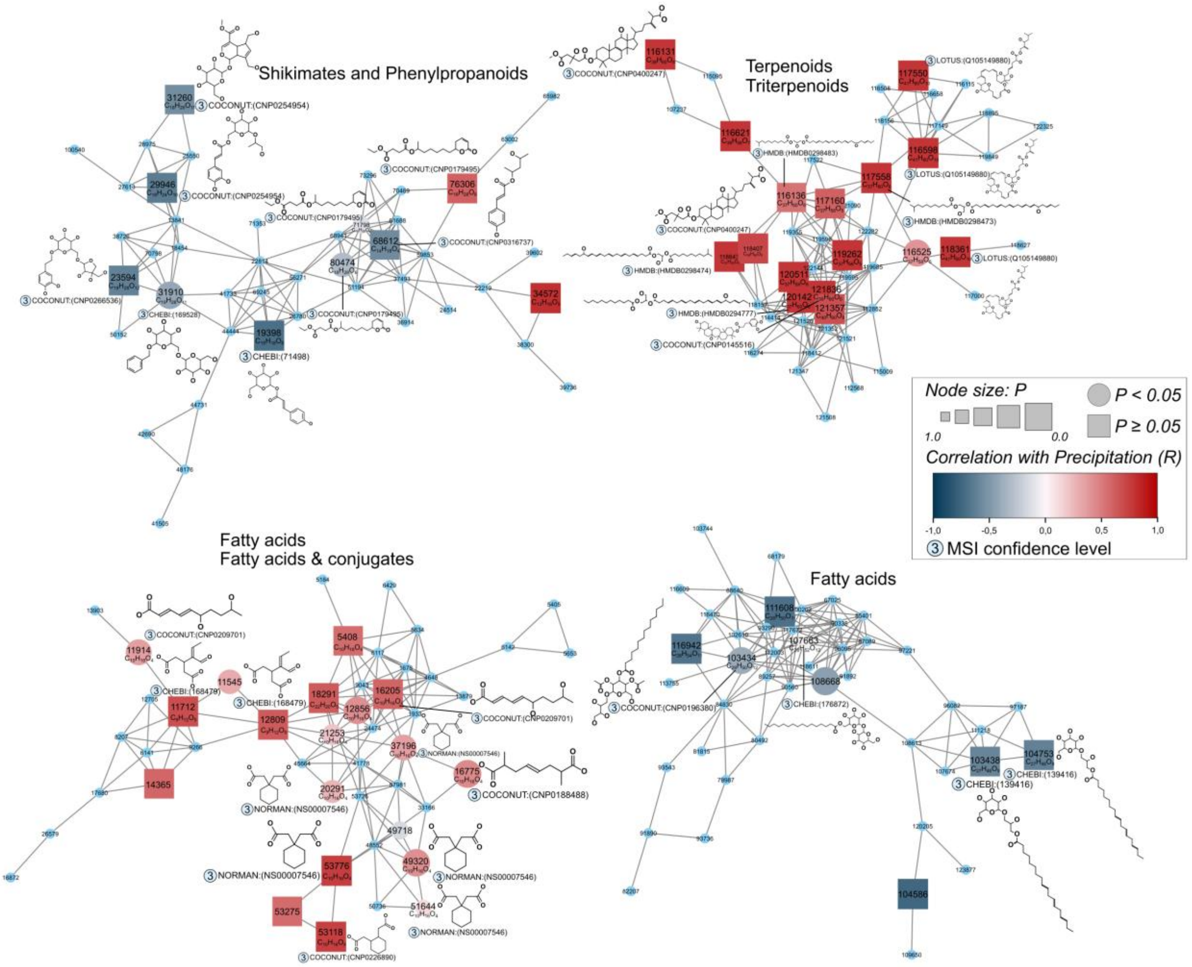


**Fig. S8 | Chemical clusters responding to precipitation.** Example of clusters of chemical features responding to precipitation anomalies. For more information, see Tables S10 and S12. Blue ellipses represent ions that did not successfully pass preprocessing. Chemical structures were drawn using Pubchem. The top 1 match from SIRIUS was used to draw the figure (MSI 2 or 3). Chemical pathways and superclasses represent the most represented ones in a given cluster. In this figure, annotations are putative and should be interpreted with caution at the compound level.

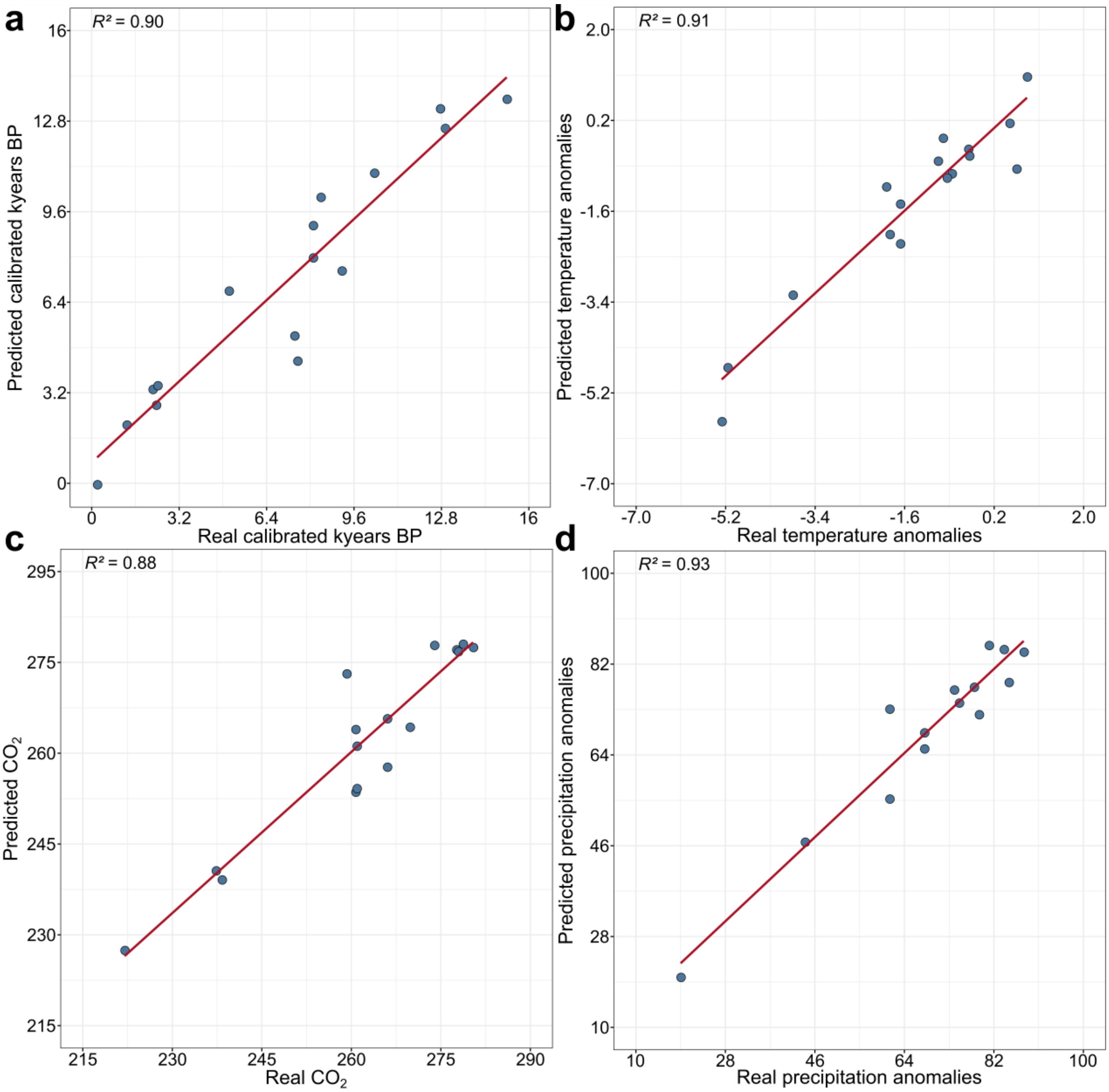


**Fig. S9 | Prediction of age and abiotic parameters using middens' metabolism. A-E.** Fits from PLSr model using significant chemical features to predict (A) the age of the midden, (B) temperature anomalies, (C) $CO_2$, and (D) precipitation anomalies. To avoid the model placing excessive weight on the two oldest samples (which inflates model performances), we removed the two oldest middens for predicting middens' age (A). A similar operation was done to predict (B) temperature anomalies and (C) $CO_2$. For all figures, the $R^2$ is related to the score of fitted predictions from the PLSr models. *kyBP: kiloyear before present.*

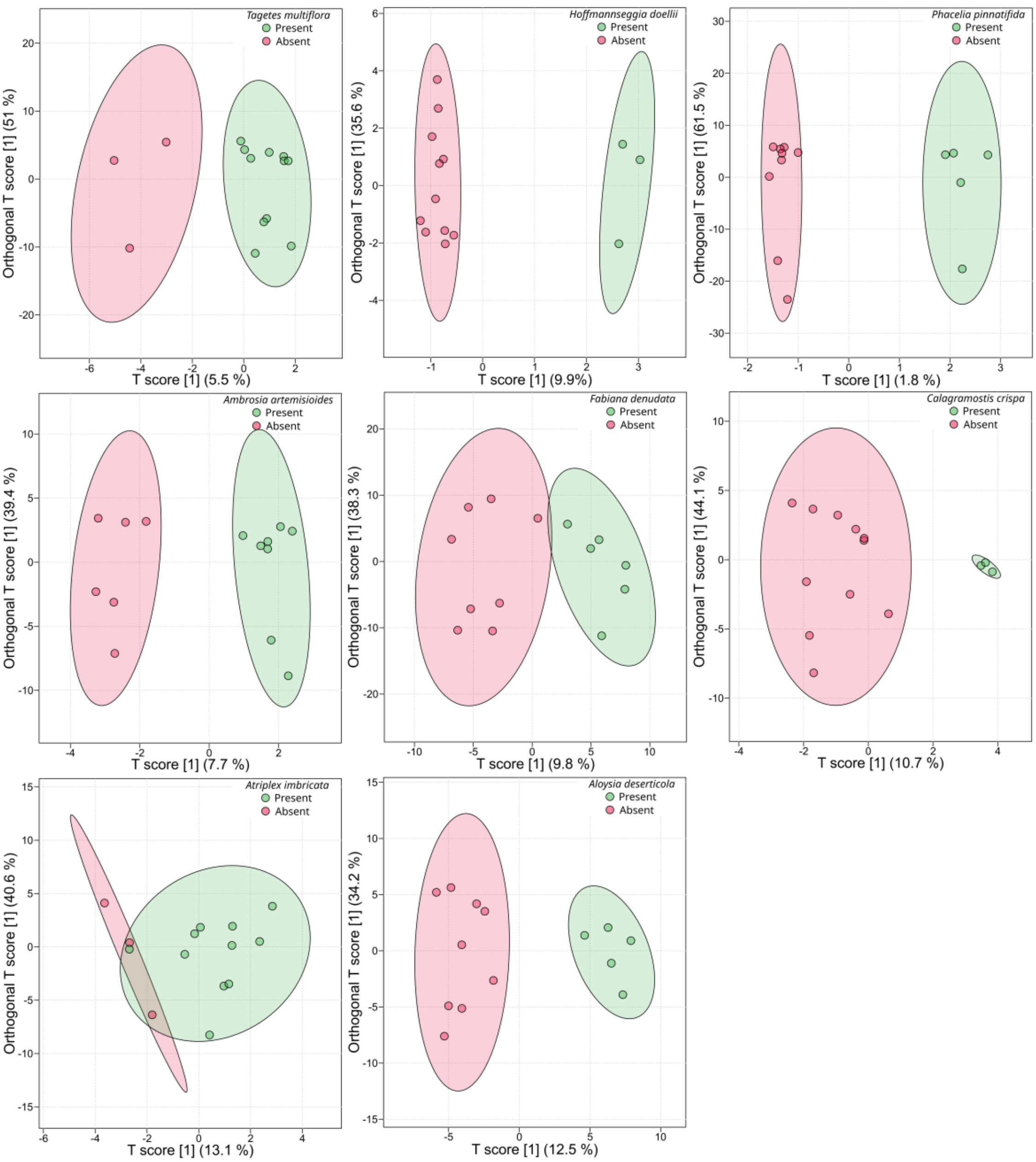


**Fig. S10 | Metabolic fingerprint can be used to predict the presence of specific plant species in the middens.** OPLS-DA of the middens in function of the presence or absence of the different plant species. Only species observed in at least three middens were used (or with at least three middens without this plant species). For each species (*e.g. Tagetes multiflora*), the metabolic features exclusive to this plant species (*i.e.* not observed in the other plant species) were first extracted and then used for the corresponding OPLS-DA.
